# Feature matrix normalization, transformation and calculation of ß-diversity in metagenomics: Theoretical and applied perspectives on your decisions

**DOI:** 10.1101/859157

**Authors:** Casper Sahl Poulsen, Frank Møller Aarestrup, Christian Brinch, Claus Thorn Ekstrøm

## Abstract

Microbial metagenomics utilising next generation sequencing is a powerful experimental approach enabling detailed and potentially complete descriptions of the microbial world around and within us. Selecting how to perform feature data normalization, transformation and calculate ß-diversity is a critical step in the analysis of metagenomic data, but also a step for which a multitude of methods are available. Researchers need to have a broad overview and understand the many methods that exist in the field and the consequences from applying them. In this perspectives article, some of the most widely used metagenomic feature data normalizations, transformations and ß-diversity metrics are discussed in the context of multivariate visualizations. We provide a framework that other researchers can utilize to evaluate how robust their test data are when applying different normalizations, transformations and ß-diversity metrics, and visually compare the results of the methods. We constructed an *in silico* test dataset to evaluate the setup and clarify how the theoretical discussion is transferable to this data. We urge other researchers to implement their own test data, normalization, transformation, ß-diversity metric and visualization methods, in the hope that it will advance better decision making both in study design and analysis strategy.

## 1 The lack of consensus on how to perform data normalization, transformation and calculate ß -diversity

Next generation sequencing (NGS) is applied heavily in microbiome research, enabling both taxonomic and functional descriptions of microbiomes (1,2). Metagenomic data need to be processed before analysis to make sample comparisons possible due to the differences in sequencing depth. Furthermore, there is an increasing awareness that such data are compositional and should be processed accordingly (3–7).

The choice of how to perform data normalization, transformation and computation of ß-diversity can have a substantial impact on the results from the subsequent data analysis especially since metagenomic data typically are sparse, because some features are not present or their abundance are below the limit of detection. Classically, metagenomic feature data are either relativized to some sample characteristics such as the number of total reads, bacterial reads etc., or from a compositional, more transparent measure according to the number of assigned reads that is also known as total sum scaling (TSS). When relativizing, the precision of measurement is lost, considering that data are heteroscedastic direct comparison of samples is flawed if methods assume equal variance. (8). Therefore, rarefying can be performed, but it has been argued against due to the loss of power (8). Relativizing is highly influenced by the most abundant features, alternatively, the median, quantile normalization or cumulative sum scaling (CSS) can be used (9,10). Methods developed for normalizing data such as trimmed mean of M-values (TMM) and relative log expression (RLE), can be relevant if most features are not changing between samples (11,12). The compositional data analysis framework provides an additional approach to analyse metagenomic data with a multitude of possibilities for estimating zeroes and visualization (13–16). There are arguably advantages and disadvantages to applying all of the different methods described (7,9,17,18).

Several R packages have implemented the techniques described above such as “vegan”, “edgeR”, “DESeq2”, “phyloseq” and “compositions” (10,13,19–21). From these packages, we have identified 228 combinations of normalizing and transforming data and calculating ß-diversity metrics. This is not an exhaustive list of possible methods to apply, and therefore processing metagenomic data is a task where tradition and ease of implementation are important factors governing researchers’ decisions. Understanding the more advanced methods, for instance, to perform compositional data analysis is most likely also a reason for these methods to not have become common as observed in other fields (22).

The aim of the present study is to provide theoretical as well as applied analytical perspectives on normalization and transformation of metagenomic data in the context of calculating ß-diversity that is used for statistical inference and multivariate visualizations. We have constructed an *in silico* dataset to visualize how data processing affects metagenomic analysis. The dataset was used to investigate if methods are robust according to sequencing depth and the influence of changes in data structure. Furthermore, a visualization of a dissimilarity matrix containing the Procrustes test results for all selected methods compared pairwise provides a comparison of how the methods resemble each other. We provided the code used to generate the analysis in the hope that other researchers can use it as a tool for assessing the effect and sensitivity of using different transformation, normalization and ß-diversity methods by incorporating their own test data, favourite methods or visualization techniques.

## 2 Theoretical perspectives on data normalization, transformation and ß-diversity calculation in metagenomics

In this section, perspectives are provided on feature data transformation, normalization and ß-diversity metrics where we, for the latter, have focused on Euclidean distance, Manhattan distance and Bray-Curtis dissimilarity due to their widespread use and acceptance in metagenomics. In terms of between sample comparisons, normalization is primarily performed to take sequencing depth into account and, transformation is performed to weigh how the differences between features should be emphasized.

One of the most common methods to account for sequencing depth is TSS. When calculating ß-diversity, this method is driven by the features with the largest differences between samples that are typically also the most abundant features because it scales linearly in absolute values. Multivariate visualisations and statistical tests therefore depend on the differences in the most abundant features. To deemphasize this effect, log transformations or square root transformations, such as the Hellinger transformation, are used. Sometimes these transformations are applied post TSS; however, if this is the case, it should be emphasized that sample values no longer add up to the same total sum anymore. If detection is the primary focus of analysis, making a presence absence (PA) transformation can be justified because this removes the effect of abundance. From a practical viewpoint, PA requires high specificity when mapping reads, commonly at the cost of sensitivity, control of contamination during sample processing and is not robust according to sequencing depth unless different detection limits are implemented prior to transformation.

Rarefying (or subsampling) provides data where precision of a measurement is the same across samples, typically performed by rarefying to the level of the sample with the fewest assigned reads, at the cost of sensitivity. The loss of precision is usually not a problem if sequencing depth is even, but a similar argument can be made in this case when relativizing. Data are still heteroscedastic and therefore should be modelled accordingly, i.e. when performing differential abundance analysis (8,9). Both relativizing and rarefying do not take the compositionality of data into account, but perform well if the most abundant features between samples are relatively constant, which is rarely known. If, on the other hand, most features are not changing between compared groups of samples, the median, RLE and TMM offer a better solution (9). This is also why RLE and TMM were implemented in DESeq2 and edgeR, respectively, for the analysis of expression data. In expression data, it is often a good assumption, for instance in a clinical study, that treatment only changes expression of a few genes (11,21). In metagenomics, this assumption could be met, but from our experience, working in the field and with spike-in organisms, this is rarely the case.

When calculating ß-diversity, the length of the straight line between two points can be calculated, this is also known as the Euclidean distance. This method would be straightforward if the points were in Euclidean space, but metagenomic data are compositional and points are therefore confined to a simplex. When calculating Euclidean distances, the differences are squared, consequently, the greatest differences are further emphasized relative to using Manhattan distance or Bray-Curtis dissimilarity. This could be counterbalanced by performing a log or Hellinger transformation. Manhattan distance is the sum of absolute differences. Manhattan distance is also the numerator of Bray-Curtis dissimilarity that is then scaled to the sum of total features in the two samples. The Manhattan distance also does not account for the compositionality of data.

## 3 Another approach to data normalization in metagenomics

A solution to the challenges described above is to use a compositional analysis framework. Using centered log ratio (CLR) transformation, where the log of each feature is compared relative to the geometric mean, or the isometric log ratio (ILR) transformation, where orthogonal basis functions are used to span the simplex space somewhat analogous to the CLR transformation, in the context of calculating ß-diversity (23,24). Performing both methods enables real-space calculations and consequently Euclidean distances when calculating ß-diversity. The methods are simple in principle, but zeroes have to be imputed and this represents a major challenge when dealing with metagenomic data that are typically sparse (4,25,26). One often-used solution is to detect features with a zero and then remove the features from all samples. This option is recommended when features are low abundant in the others samples, but in metagenomic studies, a feature might be relatively highly abundant in one sample and not present in another. Another approach is to add a pseudo-count, multiplicative simple replacement or a Bayesian approach (15,27,28). Nonetheless, in all imputations of zeroes, there is no way of knowing the difference between a “true” zero representing a feature that is not present and a zero that is below the detection limit. Imputation in this situation is therefore limited to assigning a value below the detection limit, even though the feature might not be present (27,28).

From a mathematical perspective, we expect the compositional methods to offer a desirable characteristic in that data are not constrained to the simplex, but considering sparsity, which is commonly an artefact of metagenomic data, zeroes have to be imputed.

## 4 Seeing is believing - *In silico* comparison of data normalization, transformation and ß-diversity calculation in metagenomics

To provide applied analytical perspectives, an *in silico* dataset was constructed to reflect typical challenges in metagenomic data including sparsity and differences in sequencing depth. A reference (Ref), equivalent to a sample, was created consisting of abundance profiles of 70 different organisms (i.e. number of sequence reads mapping to a given organism). The sample consisted of counts from the following abundance levels:

- **High** (1 random sampling between 1000-5000),
- **Medium high** (3 random samplings between 100-999 with replacement),
- **Medium** (9 random samplings between 5-99 with replacement),
- **Low** (27 random samplings between 0-4 with replacement), and
- **Not present** (30 zeroes).

The test data contained 70 different features (i.e. organisms), but this was a trade-off to make the analysis run on a desktop computer.

Eleven other samples were created, all variations of the reference:

- Multiplying counts with 2 (SF2) and 10 (SF10),
- Changing counts to zeroes in each of the different abundance levels (SwHato0, SwMHato0, SwMato0, SwLato0),
- Switching the highly abundant feature with one in each of the other abundance groups (SwHaMHa, SwHaMa, SwHaLa, SwHaNP), and
- Reversing the reference (RevRef) to create a very dissimilar sample only sharing a few low abundant features.

These artificial samples represent potential differences that are of interest to assess the effect of sequencing depth and structural differences in data. The full computer code documents the exact construction of the samples and their variations (https://github.com/csapou/DataProcessinginMetagenomics).

To limit the number of combinations of normalization, transformation and ß-diversity metrics in figures, we selected 36 methods. We included Euclidean distance, Manhattan distance and Bray-Curtis dissimilarity as ß-diversity metrics, since these metrics are popular in metagenomics. The selected transformation and normalization steps were based on tradition in the field of microbial ecology (TSS, rarefying, PA and CSS). We also included Hellinger and log transformation both before and after TSS. Some methods are implemented to normalize RNA-expression data (TMM and DESeq (poscount argument in estimating SizeFactors)). For methods that adhere to the compositional data analysis framework, we included six methods that use Euclidean distances. Zeroes were estimated with multiplicative simple replacement or adding a pseudo-count of one prior to TSS, then performing both CLR and ILR. We also included TSS and then added a pseudo-count of the minimum divided by ten before performing CLR and ILR.

All statistical analysis and visualization of data were performed in R version 3.4.4, and data transformation, normalization and calculation of ß-diversity were performed using the packages described above. To visualize the dissimilarities and distances between the different samples, we created a heatmap with accompanying dendrograms using complete linkage clustering of Euclidean distances based on the full-scale distance and dissimilarity matrices. Heatmaps were generated using the ‘pheatmap’ package by extracting the ß-diversity to the reference sample. ß-diversity values were made comparable in each of the methods by scaling to the max value. To compare all the distance and dissimilarity matrices pairwise, a Procrustes approach was used based on randomization tests (29,30). A dissimilarity matrix of the processing methods was created by subtracting the Procrustes correlations from one. Metric multidimensional scaling of the dissimilarity matrix was performed by running the capscale function unconstrained from ‘vegan’. The generation of the principal coordinates analysis (PCoA) plot of the first two dimensions, density plot of the correlations, stress plot containing a scatterplot of the distance observed in the PCoA as a function of the “true” ß-diversity calculated and the scree plot showing the variance in the principal components were performed with ‘ggplot2’ (31). The code to generate test data and perform data processing is provided at https://github.com/csapou/DataProcessinginMetagenomics with additional principal component analysis (PCA) plots and PCoA plots of all individual methods and randomly generated samples.

From Figure 1 we find that samples scaled by a factor of 2 or 10 had a low ß-diversity relative to the reference sample, indicating that the methods we selected were able to control the effect of sequencing depth, which is a bare minimum for applying them to this type of data. Some inconsistency was observed when performing log or log-ratio transformations. This effect can be reduced in this case by estimating zeroes at a lower level. The reverse sample representing a dissimilar community was also generally the one with the highest ß-diversity relative to the reference. ß-diversity metrics generally cluster containing either Euclidean distances or Bray-Curtis dissimilarity together with Manhattan distance. Bray-Curtis dissimilarity and Manhattan distance cluster when performing TSS, because the denominator evaluates to 2 when calculating the Bray-Curtis dissimilarity and is therefore just a factor of two scaling of the Manhattan distance. Transformation and normalization methods also cluster to some extent.

**Figure 1:**
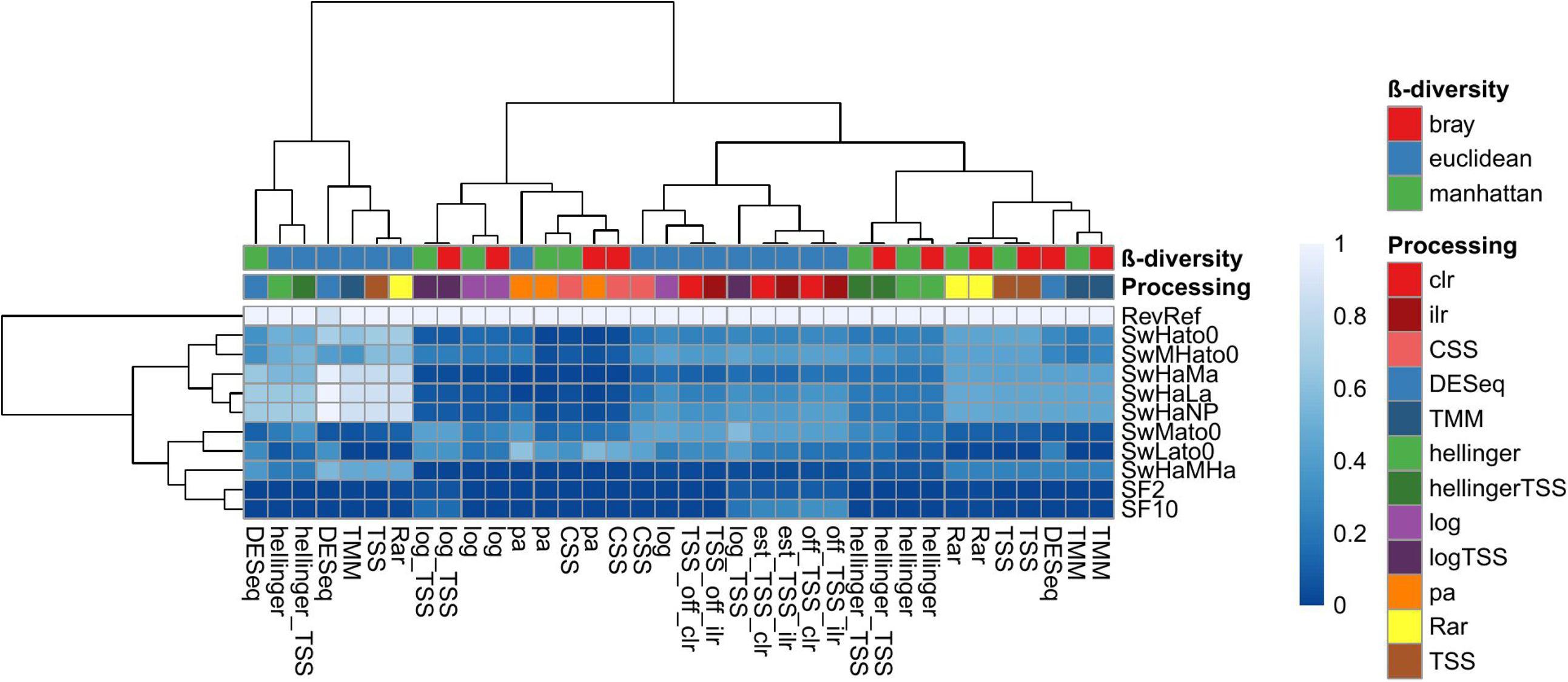
Heatmap visualizing the ß-diversity to the reference sample using different strategies to normalize and transform data. The ß-diversity relative to the reference for each method was normalized according to the max value. Dendograms were created using complete linkage clustering of Euclidean distances. The rows in the heatmap represent different modifications to the reference, where RevRef represents reversing the reference, Sw represents switching, Ha represents high abundance, MHa represents medium high abundance, Ma represents medium abundance, La represents low abundance, NP represents not present, and SF represents scaling factor. The column labels in the heatmap contain extended explanations of zero estimation, where TSS represents total sum scaling, Rar represents rarefying, pa represents presence absence, CSS represents cumulative sum scaling, off represents an offset of zeroes, est represents a zero estimate using multiplicative simple replacement, ilr represents isometric log ratio transformation, clr represents centred log ratio transformation, and TMM represents trimmed mean of M-values.

In Figure 2, where the processing methods are compared pairwise in full scale, the classical methods show a spectrum between abundance-driven processing exemplified by TSS and PA (Fig. 2A). In between these extremes, variations of Hellinger and log transformations are plotted as we expected from the theoretical discussion. The methods adhering to the compositional data analysis framework do not cluster, emphasizing the need for further investigations into the effect of estimating zeroes. The methods developed for normalization of gene expression data to perform differential abundance analysis are likely to perform badly with this *in silico* data because they assume that large proportions of features are constant between samples. Comparing communities that are highly different, for example, the reverse sample in this dataset makes these methods inappropriate. The validation plots in the form of stress plot and scree plot show that the observed dissimilarity correlates with the ordination distance, and a large proportion of the variance is explained in the first two axes, respectively (Fig. 2C-D).

**Figure 2:**
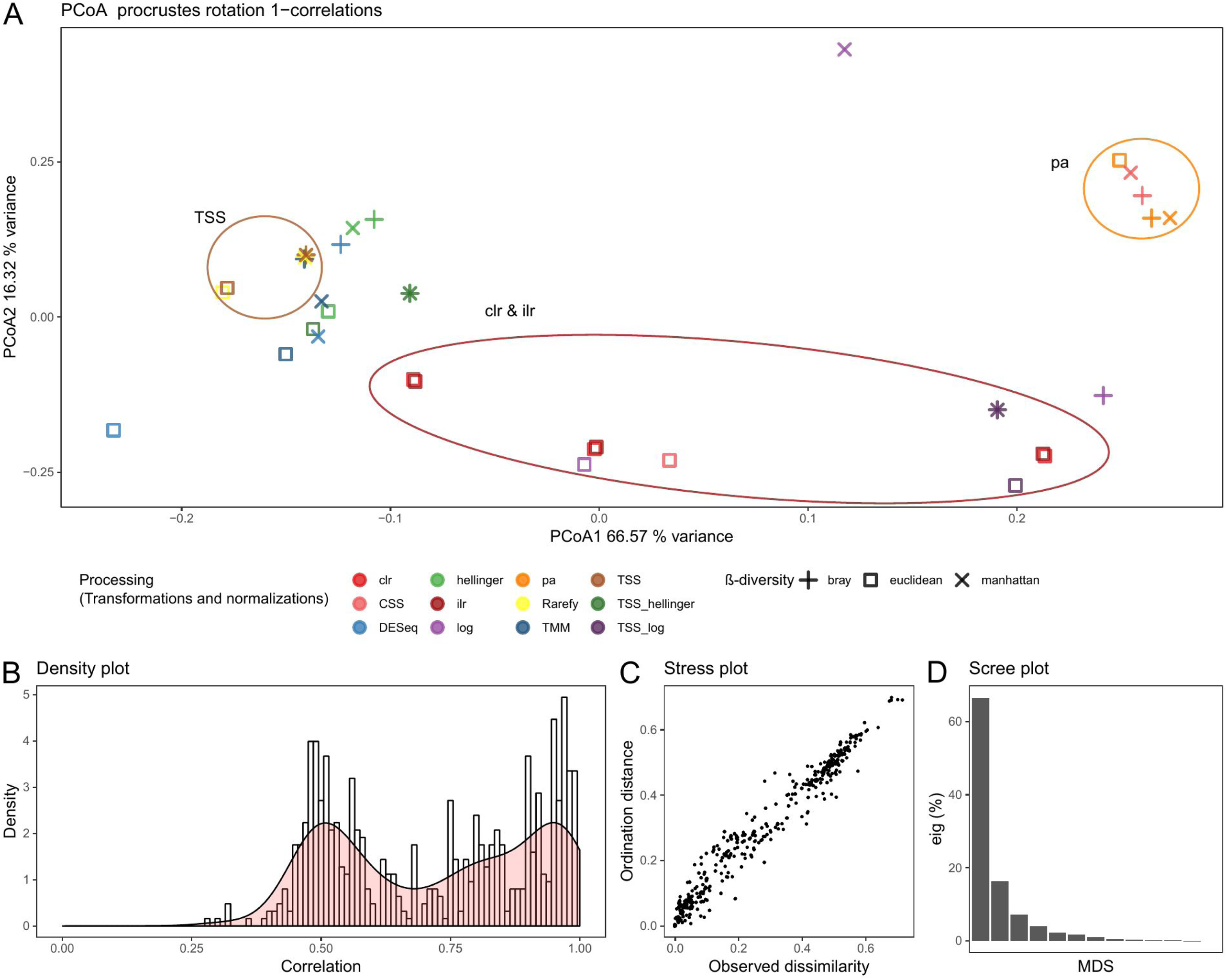
A: Principal coordinates analysis (PCoA) of the dissimilarity matrix containing pairwise comparisons of 1 - Procrustes correlations between methods, B: Density plot of correlations, C: Stress plot comparing observed dissimilarity to ordination distance in the PCoA and D: Scree plot of the percent of variation explained by the axes. A dissimilarity matrix was created from all of the pairwise comparisons of metagenomics data analysis pipelines represented by one minus the Procrustes correlation. Redundancy analyses were performed unconstrained using the capscale function in vegan creating PCoA-, density, stress- and scree plot with ggplot2. Ellipses where added manually highlighting presence absence (pa), total sum scaling (TSS), and the compositional methods centred log ratio transformation (clr) and isometric log ratio transformation (ilr). In the processing (transformations and normalizations) legend, CSS represents cumulative sum scaling, and TMM represents trimmed mean of M-values. A small amount of jitter was added to distinguish clr and ilr.

## 5 Multivariate visualization in metagenomics - a step forward

We hope that the theoretical perspectives together with the visualizations provided demonstrate that data normalization, transformation and calculation of ß-diversity have a substantial impact on the analysis and multivariate visualization of metagenomic data. We consider the public source code as a resource that other researchers can utilize to implement their own favourite methods for processing metagenomic data (https://github.com/csapou/DataProcessinginMetagenomics). Here, we also provide all of the 228 methods that we have identified with additional randomly generated samples. To perform a sensitivity analysis of the effect of using different data normalization and transformation strategies in the context of calculating ß-diversity, a density plot is provided for all of the Procrustes test correlations. From the analysis on our test dataset we see that there are two peaks with approximately the same height and the lower one is centred around a correlation of 0.5, indicating that data processing is important for this test data (Fig. 2B). On the other hand, performing this analysis on another test dataset might reveal high correlations between all methods. This would indicate that the conclusions derived from the data are robust to withstand applying different normalizations, transformations and ß-diversity metrics.

Other relevant modifications include the removal of the reversed sample from the analysis to look at the subtle differences between similar samples. With the large number of combinations of normalizations, transformations and ß-diversity metrics to select from, we discourage other researchers from implementing their real data to circumvent pipeline-hacking analogous to p-hacking (32). A better option for the users would be to implement their own relevant test dataset, and from this analysis, and together with theoretical considerations, select one or a few processing methods before analysing their real data. We hope that the code provided also eases the implementation of new methods. Generation of dendograms in the heatmap and the PCoA of Procrustes test results were run using default settings, and an investigation could also be initiated to assess how this might influence the results. Again, we urge others to implement their own favourite methods.

We would like to highlight other aspects of good scientific practice in metagenomics and refer readers to articles on study design (33–35), sample processing (36–39) and other aspects of metagenomic data analysis primarily focusing on differential abundance analysis of features (3,7–9,40–42).

## 6 Acknowledgments

The authors wish to thank Jeffrey Skiby for language editing.

## 7 Author Contributions Statement

CP and CE conceived the ideas and wrote the paper. CP analysed the data, made the figures and performed the literature review. FA and CB revised the manuscript. All authors read and approved the manuscript.

## 1 Data Availability Statement

The datasets generated and analyzed for this study can be found at: https://github.com/csapou/DataProcessinginMetagenomics

## References

1. Kim M, Lee K-H, Yoon S-W, Kim B-S, Chun J, Yi H. Analytical Tools and Databases for Metagenomics in the Next-Generation Sequencing Era. Genomics Inform [Internet]. 2013;11(3):102. Available from: https://synapse.koreamed.org/DOIx.php?id=10.5808/GI.2013.11.3.102

2. Land M, Hauser L, Jun S-R, Nookaew I, Leuze MR, Ahn T-H, et al. Insights from 20 years of bacterial genome sequencing. Funct Integr Genomics. 2015;

3. Nayfach S, Pollard KS. Toward Accurate and Quantitative Comparative Metagenomics. Cell [Internet]. 2016;166(5):1103–16. Available from: http://dx.doi.org/10.1016/j.cell.2016.08.007

4. Gloor GB, Reid G. Compositional analysis: a valid approach to analyze microbiome high-throughput sequencing data. Can J Microbiol. 2016;62(8):692–703.

5. Gloor GB, Macklaim JM, Pawlowsky-Glahn V, Egozcue JJ. Microbiome datasets are compositional: And this is not optional. Front Microbiol. 2017;8:1–6.

6. Odintsova V, Tyakht A, Alexeev D. Guidelines to Statistical Analysis of Microbial Composition Data Inferred from Metagenomic Sequencing. Curr Issues Mol Biol [Internet]. 2017;24:17–36. Available from: http://www.caister.com/cimb/abstracts/v24/17.html

7. Weiss S, Xu ZZ, Peddada S, Amir A, Bittinger K, Gonzalez A, et al. Normalization and microbial differential abundance strategies depend upon data characteristics. Microbiome. 2017;5(1):1–18.

8. McMurdie PJ, Holmes S. Waste Not, Want Not: Why Rarefying Microbiome Data Is Inadmissible. PLoS Comput Biol. 2014;10(4).

9. Pereira MB, Wallroth M, Jonsson V, Kristiansson E. Comparison of normalization methods for the analysis of metagenomic gene abundance data. BMC Genomics. 2018;19(1):1–17.

10. Oshlack A, Robinson MD. A scaling normalization method for differential expression analysis of RNA-seq data. Genome Biol. 2010;11.

11. Robinson MD, Oshlack A. A scaling normalization method for differential expression analysis of RNA-seq data. Genome Biol. 2010;11(R25).

12. Love MI, Huber W, Anders S. Moderated estimation of fold change and dispersion for RNA-seq data with DESeq2. Genome Biol. 2014;15(550).

13. Boogaart AKG Van Den, Tolosana-delgado R, Bren M, Boogaart MKG Van Den. Package ‘compositions.’ 2018;

14. Cao KAL, Costello ME, Lakis VA, Bartolo F, Chua XY, Brazeilles R, et al. MixMC: A multivariate statistical framework to gain insight into microbial communities. PLoS One. 2016;

15. Palarea-Albaladejo J, Martín-Fernández JA. ZCompositions - R package for multivariate imputation of left-censored data under a compositional approach. Chemom Intell Lab Syst [Internet]. 2015;143:85–96. Available from: http://dx.doi.org/10.1016/j.chemolab.2015.02.019

16. Silverman JD, Washburne AD, Mukherjee S, David LA. A phylogenetic transform enhances analysis of compositional microbiota data. Elife. 2017;6:1–20.

17. Dillies MA, Rau A, Aubert J, Hennequet-Antier C, Jeanmougin M, Servant N, et al. A comprehensive evaluation of normalization methods for Illumina high-throughput RNA sequencing data analysis. Brief Bioinform. 2013;14(6):671–83.

18. Parks DH, Beiko RG. Measures of phylogenetic differentiation provide robust and complementary insights into microbial communities. ISME J [Internet]. 2013;7(1):173–83. Available from: http://dx.doi.org/10.1038/ismej.2012.88

19. Oksanen J et al. vegan: Community Ecology Package [Internet]. 2019. Available from: https://cran.r-project.org/package=vegan

20. McMurdie PJ, Holmes S. Phyloseq: An R Package for Reproducible Interactive Analysis and Graphics of Microbiome Census Data. PLoS One. 2013;

21. Anders S, Huber W. Differential expression analysis for sequence count data. Genome Biol. 2010;11(R106).

22. Filzmoser P, Hron K, Reimann C. The bivariate statistical analysis of environmental (compositional) data. Sci Total Environ [Internet]. 2010;408(19):4230–8. Available from: http://dx.doi.org/10.1016/j.scitotenv.2010.05.011

23. Aitchison J. The Statistical Analysis of Compositional Data. London: Chapman and Hall; 1986.

24. Egozcue J, Pawlowsky Glahn V, Mateu-Figueras G, Barceló Vidal C. Isometric logratio for compositional data analysis. Math Geol. 2003;35(3):279–300.

25. Costea PI, Zeller G, Sunagawa S, Bork P. A fair comparison. Nat Methods. 2014;11(4):359.

26. Tsilimigras MCB, Fodor AA. Annals of Epidemiology Compositional data analysis of the microbiome : fundamentals, tools, and challenges. Ann Epidemiol [Internet]. 2016;26(5):330–5. Available from: http://dx.doi.org/10.1016/j.annepidem.2016.03.002

27. Martín-Fernández JA, Barceló-Vidal C, Pawlowsky-Glahn V. Dealing with Zeros and Missing Values in Compositional Data Sets Using Nonparametric Imputation. Math Geol. 2003;35(3):253–78.

28. Martín-Fernández JA, Hron K, Templ M, Filzmoser P, Palarea-Albaladejo J. Bayesian-multiplicative treatment of count zeros in compositional data sets. Stat Modelling. 2015;15(2):134–58.

29. Jackson D. PROTEST: A PROcrustean Randomization TEST of community environment concordance. Ecoscience. 1995;2(3):297–303.

30. Peres-Neto PR, Jackson DA. How well do multivariate data sets match? The advantages of a procrustean superimposition approach over the Mantel test. Oecologia. 2001;129(2):169–78.

31. Wickham H. ggplot2: Elegant Graphics for Data Analysis. New York: Springer-Verlag; 2016.

32. Simonsohn U, Nelson LD, Simmons JP. P-curve: A key to the file-drawer. J Exp Psychol Gen. 2014;143(2):534–47.

33. Prosser JI. Replicate or lie. Environ Microbiol. 2010;12(7):1806–10.

34. Lennon JT. Replication, lies and lesser-known truths regarding experimental design in environmental microbiology. Environ Microbiol. 2011;13(6):1383–6.

35. en Hoopen P, Finn RD, Bongo LA, Corre E, Fosso B, Meyer F, et al. The metagenomic data life-cycle: Standards and best practices. Gigascience. 2017;6(8):1–11.

36. Costea PI, Zeller G, Sunagawa S, Pelletier E, Alberti A, Levenez F, et al. Towards standards for human fecal sample processing in metagenomic studies. Nat Biotechnol [Internet]. 2017;35(11):1069–76. Available from: http://dx.doi.org/10.1038/nbt.3960

37. Wiehlmann L, Pienkowska K, Hedtfeld S, Dorda M, Tümmler B. Impact of sample processing on human airways microbial metagenomes. J Biotechnol [Internet]. 2017 May 20 [cited 2018 Jan 2];250:51–5. Available from: http://linkinghub.elsevier.com/retrieve/pii/S0168165617300123

38. Jones MB, Highlander SK, Anderson EL, Li W, Dayrit M, Klitgord N, et al. Library preparation methodology can influence genomic and functional predictions in human microbiome research. Proc Natl Acad Sci U S A [Internet]. 2015 Nov 10 [cited 2018 Jan 2];112(45):14024–9. Available from: http://www.pnas.org/lookup/doi/10.1073/pnas.1519288112

39. Knudsen BE, Bergmark L, Munk P, Lukjancenko O, Priemé A, Aarestrup FM, et al. Impact of Sample Type and DNA Isolation Procedure on Genomic Inference of Microbiome Composition. Jansson JK, editor. mSystems [Internet]. 2016 Oct 25 [cited 2018 Jan 2];1(5):e00095–16. Available from: http://msystems.asm.org/lookup/doi/10.1128/mSystems.00095-16

40. Jonsson V, Österlund T, Nerman O, Kristiansson E. Statistical evaluation of methods for identification of differentially abundant genes in comparative metagenomics. BMC Genomics [Internet]. 2016;17(1):1–14. Available from: http://dx.doi.org/10.1186/s12864-016-2386-y

41. Thorsen J, Brejnrod A, Mortensen M, Rasmussen MA, Stokholm J, Al-Soud WA, et al. Large-scale benchmarking reveals false discoveries and count transformation sensitivity in 16S rRNA gene amplicon data analysis methods used in microbiome studies. Microbiome [Internet]. 2016 Dec 25 [cited 2019 Nov 28];4(1):62. Available from: http://microbiomejournal.biomedcentral.com/articles/10.1186/s40168-016-0208-8

42. Russel J, Thorsen J, Brejnrod AD, Bisgaard H, Sørensen SJ, Burmølle M. DAtest: a framework for choosing differential abundance or expression method. bioRxiv [Internet]. 2018 Jan 2 [cited 2019 Nov 28];241802. Available from: https://www.biorxiv.org/content/10.1101/241802v1

